# AQuA2-Cloud: a web platform for fluorescence bioimaging activity analysis

**DOI:** 10.64898/2026.03.06.709938

**Authors:** Mark Bright, Xuelong Mi, Daniela Duarte, Erin Carey, Boyu Lyu, Yihzi Wang, Axel Nimmerjahn, Guoqiang Yu

## Abstract

**Background:** Advanced biological imaging analysis platforms such as Activity Quantification and Analysis (AQuA2) enable accurate spatiotemporal activity analysis across diverse cell populations within many species. These tools are increasingly important for investigating cellular signaling dynamics and behavior. However, despite advances in the accuracy and species capability of AQuA2, it remains computationally demanding for analysis of long time-series datasets and requires all users to maintain a MATLAB license, which may limit accessibility and large-scale deployment.

**Results:** To address these limitations, we have designed and made available *AQuA2-Cloud*, a portable software stack and web platform developed as an improvement and further evolution of AQuA2. This container-deployable system permits multi-user cloud-based high accuracy activity quantification with intuitive workflows, export of analysis data and project files, and comparable processing times. The platform offers integrated features such as in-browser analysis control interfaces, asynchronous program state control, multiple users and user management, support for unreliable connections, file uploading and downloading via web browsers and File Transfer Protocol, and centralized organization of analysis output.

**Conclusion:** AQuA2-Cloud constitutes a cloud-native solution for laboratories or research groups seeking to centralize analysis of spatiotemporal biological imaging datasets while reducing software installation and licensing barriers for end users. The platform enables researchers with minimal technical expertise to perform advanced bioimaging analysis through standard web browsers while maintaining the analytical capabilities of AQuA2. AQuA2-Cloud source code, deployment procedures, and documentation are freely available at (https://github.com/yu-lab-vt/AQuA2-Cloud).

## Background

Spatiotemporal biological imaging data in which various indicators provide quantifiable information pertinent to the cellular and molecular activity across the whole body allows investigators to interrogate and study behavioral and signaling dynamics [1-5]. Such data can offer discernible patterns between molecular activity and biological functionality of cells captured within an optical field or volume, and significant recent work has been conducted to obtain novel imaging data from unique perspectives and under differing modalities [6-7]. Molecular sensors are a popular mechanism to provide analyzable time-series signals during cell activity [1-3, 8-9]. Various features of detected signals such as size, duration, propagation, and spatiotemporal patterns relative to other signals can yield rich insight into the biological significance of such activity.

A variety of bioimaging data processing tools focused on spatiotemporal fluorescence activity analysis have been developed in multiple languages and various architectures and are usable across several execution environments [10-15]. However, investigators seeking to utilize these tools must potentially navigate prohibitive aspects concerning installation, configuration, and usage. CaImAn and Suite2p require user familiarity with Python environments [10-11]. AQuA, CalciumCV, and Begonia all require the end user to have a license of MATLAB [12-14]. These tools can be computationally expensive for some analyses, require specific operating environments, or expect users to have technical experience with software installation and configuration or command line usage [10-12, 16].

A novel tool called Activity Quantification and Analysis (AQuA2) was recently developed as an advanced successor to Astrocyte Quantification and Analysis (AQuA) and provides superior analytical performance when compared to its predecessor and peer methods [12, 16]. As an event-based method, AQuA2 features improved versatility and applicability, and incorporates a new top-down analysis approach that more effectively utilizes information from across the entire optical field. Usage of Bi-directional pushing with Linear Component Operations (BILCO), an innovative machine learning algorithm, in combination with the new top-down approach enables improved accuracy and efficiency [17]. These improvements were made in response to growing usage and popularity of AQuA as well as growing complexity and size of source datasets. However, AQuA2 still requires all users to retain a license for MATLAB and all tools referenced so far conduct computations locally.

We present a cloud-based system called AQuA2-Cloud (**Figure 1**) to allow spatiotemporal bioimaging data analysis in a software-as-a-service (SaaS) format through a web-browser accessible server with an intuitive user workflow (**Figure 2**) while retaining comparable processing times and accuracy with respect to the newest tool, AQuA2. The system is written using a combination of Javascript, PHP and MATLAB, and is packaged to be flexibly deployable in a variety of environments and hardware. A web browser is used to interact with the system website which provides areas for data management, application instance control, and conduct of analyses. After a one-time deployment on a host platform, this system allows end users with minimal to no technical knowledge the ability to now perform such analysis with only a web browser and a File Transfer Protocol (FTP) client without any license requirements or environment restrictions.

**Figure 1:**
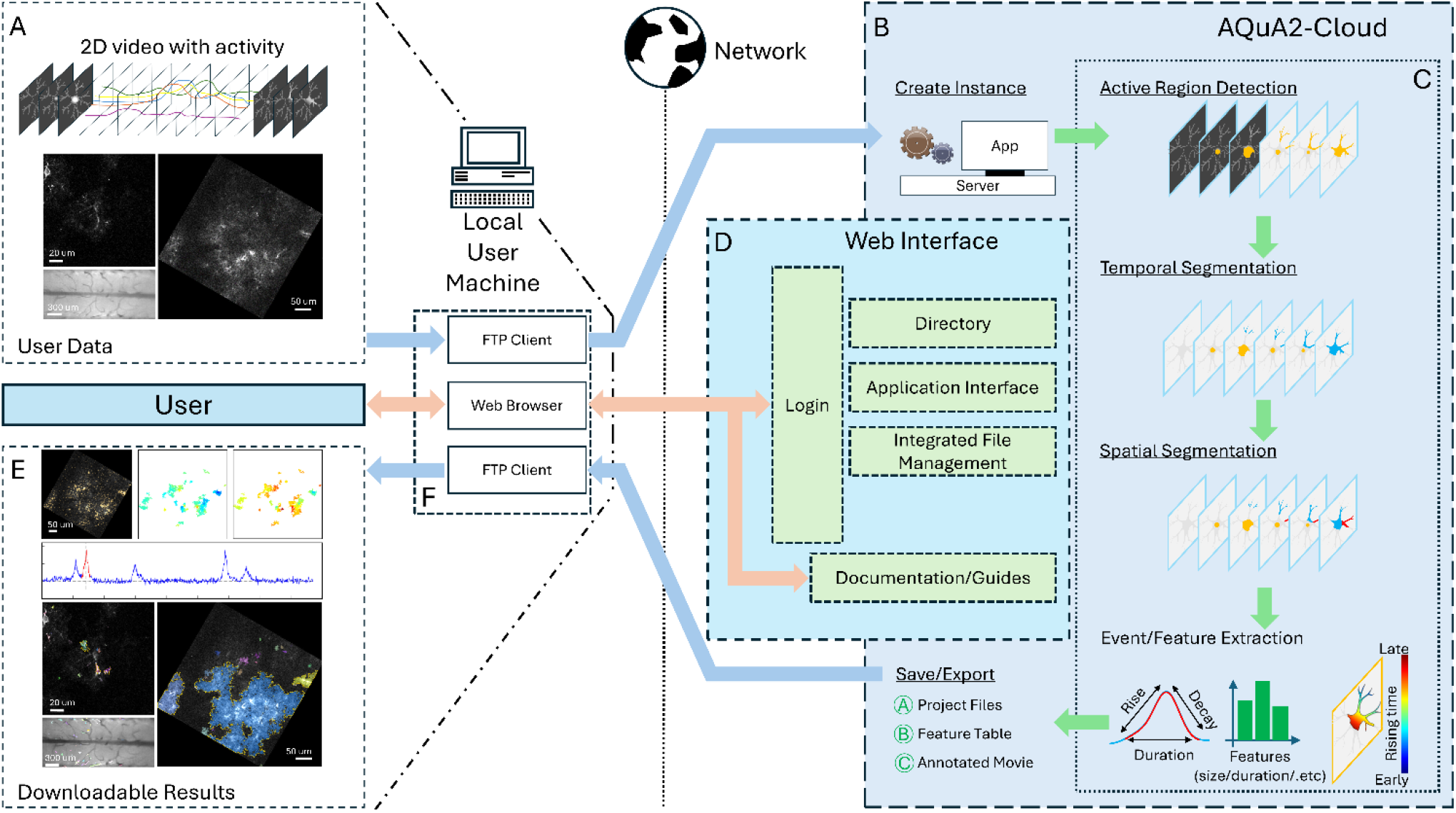
AQuA2-Cloud workflow. (A) Compatible and analyzable input data consists of 2 spatial dimension imaging movies with time as the third dimension and in which intensity-based and grouped pixel activity is present. Three exhibit imaging movies are depicted; *ex vivo* Ca^2+^ (A)(Top-left), *in vivo* Ca^2+^ (A)(Right), and mouse spinal cord Ca^2+^ (A)(Bottom-left). (B) AQuA2-Cloud functional system utilizing AQuA2’s event-based top-down signal detection and core processing pipeline (C) with an (D) asynchronous networked browser-based control interface. Analysis results (E) consisting of annotated imaging movies, detected event details, project files, and reported features are displayed in-browser and are downloadable via the web-browser and via File Transfer Protocol (F).

**Figure 2:**
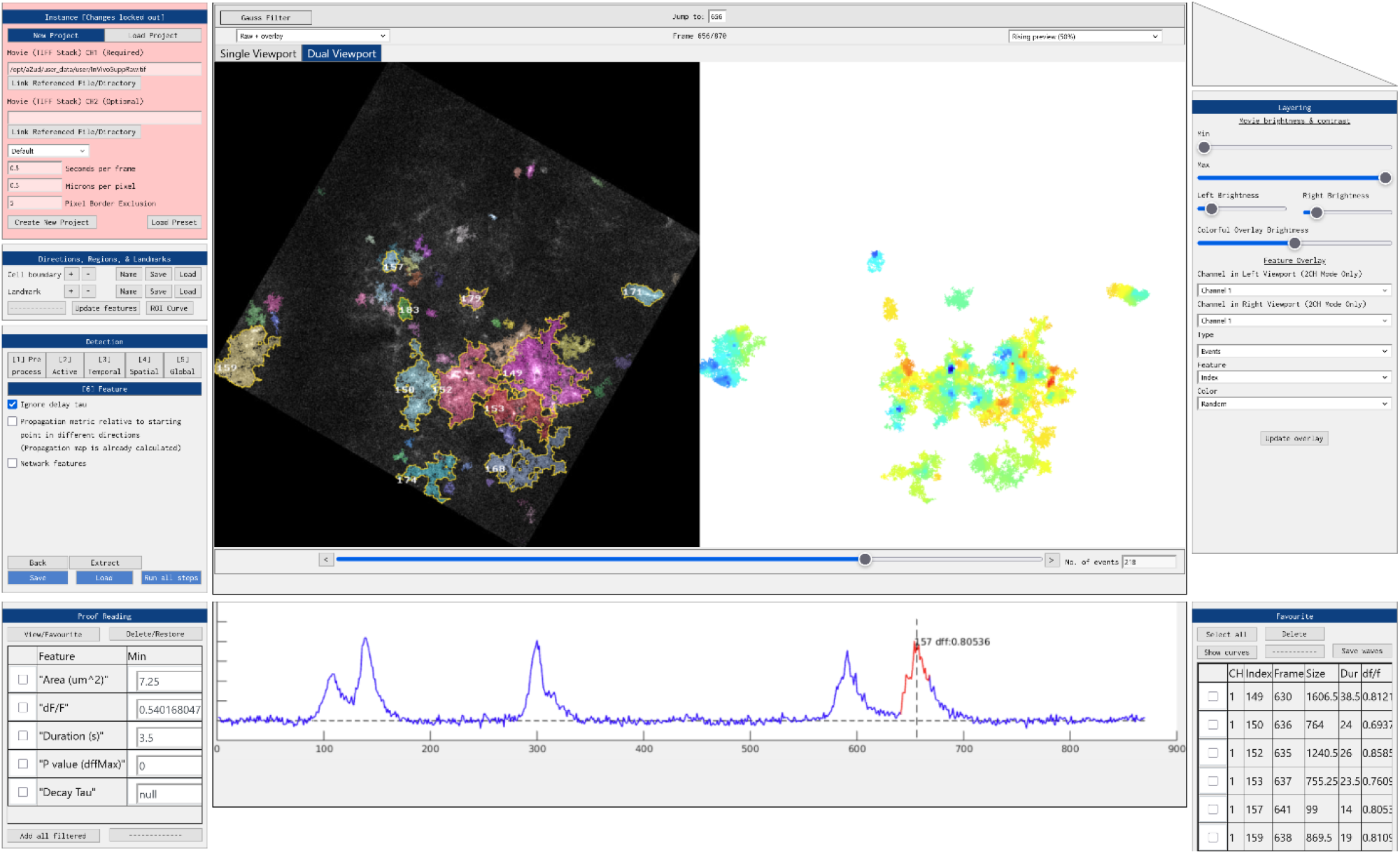
AQuA2-Cloud web interface. The web interface is designed for intuitive use, with parameters and controls grouped into panels according to functionality. End users can conduct interactive spatiotemporal activity analyses and obtain intermediate and final results comparable to those generated by AQuA2, as well as observe relevant features of interest for selected events. For each detected event, signal curve plots and key quantitative metrics are provided, including event size, propagation, duration, and signal intensity. Panel background colors change to indicate the interactability of controls and parameters given the current program state. Event filtering by user-defined criteria and frame-by-frame inspection of analysis results across the full movie are supported.

Users provide 2D spatial biological imaging data (**Figure 1A**) that is uploaded to the deployed service via their local machine (**Figure 1F**), after which they can start an AQuA2-Cloud logical MATLAB instance (**Figure 1B**). Once initialization of this logical instance is complete, it will be connected to the web browser of the user (**Figure 1D**) such that they are then able to visually and interactively process their data (**Figure 1C**). After all workflow parameter tuning and analyses, result data can be downloaded to their machine such as imaging movies with pixel-wise superimposed detected activity, event curves, and signal propagation maps (**Figure 1E**). Thus, users can now offload all expensive computation to a remote server while still being able to intuitively interact with all stages of the analysis workflow. We discuss further technical details of the operation and implementation of this system, as well as verification activity with analyses affirming compute time equivalency and correctness of cell activity signal detection.

### Implementation

AQuA2-Cloud builds on the algorithms and analysis pipeline within AQuA2 and utilizes portions of its MATLAB code for the system logical back-end. Using an Apache web server serving JavaScript pages and running PHP code, it provides a networked and lightweight user interface system and associated control mechanisms to enable remote users to perform highly configurable imaging analysis entirely from their local machines. The layout and design of user interfaces within the analysis portions of the system are visually and functionally similar to that of AQuA2 with regards to the processing workflow and the detection pipeline. A custom client-server architecture (**Figure 3**) provides capability for simultaneous parallel instances instantiated by multiple users via an integrated authentication system complete with asynchronous program state control and thus support for unstable connections inherently by design [18-22]. Users enter the URL or network address of the hosting server to arrive at a login page on the system’s website. The deployer of the system can elect to create individual user accounts for persons expected to use the system, or create public accounts and share credentials freely. After login, end users are automatically redirected to a landing page containing information needed to connect and transfer data via FTP, and a directory allowing access to guides, the integrated file manager, and the main analysis and workflow page.

**Figure 3:**
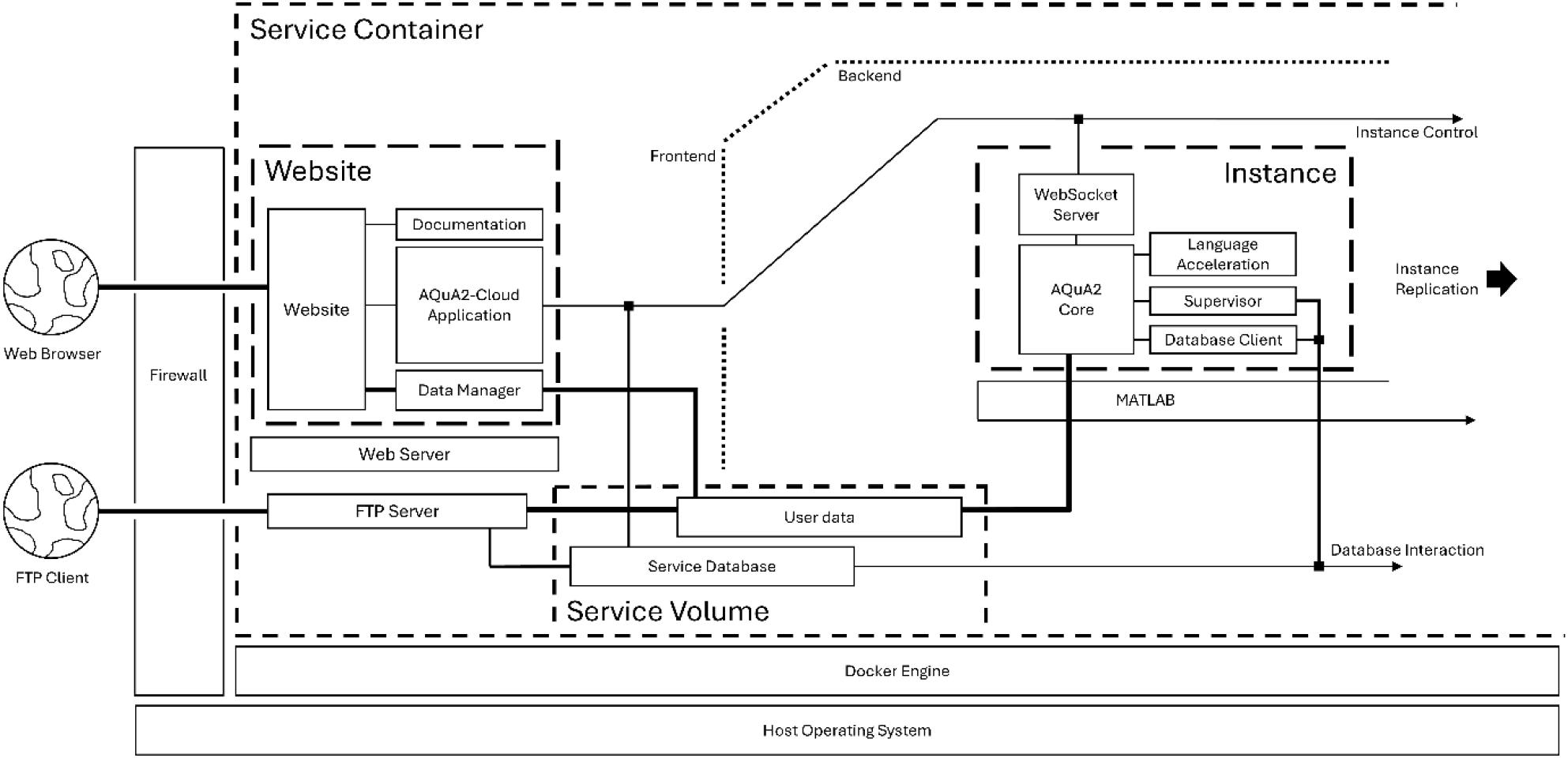
General system architecture. Depiction of functional data-flow between system components. The system website, application logic, database service, internal communication layer, and File Transfer Protocol server all run within a self-contained docker container. Users access the cloud-based application through a web browser, while large data transfers are supported via File Transfer Protocol clients.

Once raw data and any configuration files have been prepared and uploaded, users start a logical instance of the MATLAB back-end code, with this back-end instance then automatically connecting via a WebSocket system to the user’s session within the Apache web server. The analysis session’s monitored state and tracking of user logical instances is supported by MySQL, a relational database management system (RDBMS). Asynchronous program state control is made possible through having the client web session periodically poll database tables via JavaScript and PHP for the user logical session to track logical instance status information. Depending on the information reported in this database lookup, the JavaScript logic on the remote user machine modifies its behavior accordingly. This subsystem partly addresses the fundamentally unreliable nature of remote networked connections between the user and server regardless of whether the access route is across a Local Area Network (LAN) or a Wide Area Network (WAN) in that it allows users to connect to an unaffected back-end logical instance in cases where they unexpectedly disconnect or lose connection. Infrastructure and support for uploading, downloading, and organizational control of imaging datasets as well as configuration files is enabled by combined integration of a Very Secure FTP Daemon (VSFTPD) server and a PHP-based in-browser file manager. The in-browser file manager provides uploading capability for smaller files, and no size limit exists for downloading files via this file manager nor when uploading or downloading files via the integrated VSFTPD server. User credentials for general cloud-service access as well as FTP have been homogenized within the RDBMS system, further enhancing ease of use and data transfer. User management via database table modifications is possible via the deployed system’s docker container command line interface (CLI) and preprepared documentation describing the commands and procedures to be used.

One of the challenges with a cloud-based imaging data processing application is the streaming of imagery frame data to clients. An effective workflow allows users to immediately see changes resulting from manipulation of analysis parameters. The AQuA2-Cloud application page uniquely offers in-browser based mechanisms for specifying workflow parameters in an interactive environment with step-wise visual feedback (**Figure 4**) and dynamic streaming of this data throughout the entirety of the process. When users enact a parameter change, new still-frame imagery is sent to the client depending on whether the alteration calls for it. Therefore, the lightweight architecture, developed with dynamic image frame transmission as a cornerstone, allows the key feature of displaying updated imagery frames as users progress through the pipeline at various down-sampled resolutions as part of low-bandwidth mitigation. Another challenge with cloud-based applications are cases where the end user connection is maintained but experiences a high degree of packet loss or unreliability. At the architectural level, this issue is managed through the implementation of aperiodic control commands and state synchronization mechanisms within the JavaScript code and Apache, enabling bidirectional data transmission between the system and the corresponding user instance to be dynamically deferred until the connection stabilizes. The combination of Relational Database Management System querying, system routines aware of connection quality, and still-frame imagery downsampling all satisfactorily address the challenges of unreliable or dropped user connections across arbitrary network types. The architecture has also been designed with a focus on security as the integrated MySQL RDBMS is utilized for storing securely hashed authentication credentials while the cloud-service website employs HTTPS. Server-sided cryptographic functions provide secure access control checks before allowing users access to both instantiated instances as well as in-browser data storage management functions. Furthermore, running instances are tracked and checked for their status, such as for inactivity, through periodic query of the database system as part of implemented resource management and abandoned instance cleanup.

**Figure 4:**
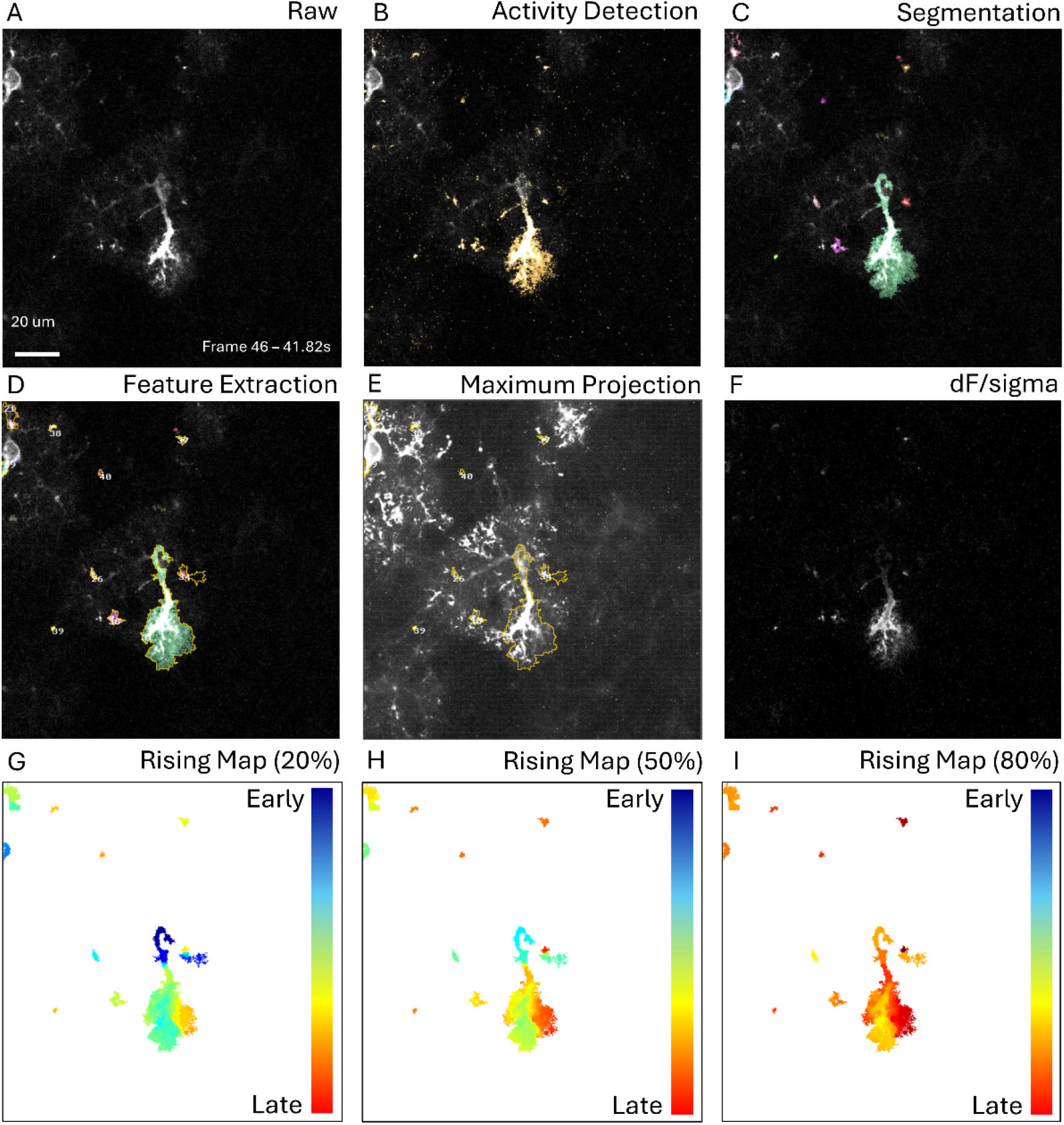
Detection pipeline step-wise results for *ex vivo* data. The analysis interface of AQuA2-Cloud is such that users can execute processing stages and obtain intermediate and final results similarly to AQuA2; as well as observe relevant features of interest. For a given raw frame (A), monitoring of detected active voxels (B) allows adjustment of parameters such that diverse signals can be captured then segmented (C) to allow eventual feature extraction (D). After detection pipeline execution, metrics evaluation for maximal event territory (E), dynamic fluorescence (F), and detected event temporal propagation (G-I) on a frame-by-frame basis are immediately viewable.

## Results

To demonstrate the efficacy and compute time equivalency of this system, we performed spatiotemporal activity analysis on three imaging movie datasets containing fluorescence activity yielded by genetically encoded molecular sensors which consists of *in vivo* (2 Hz) optical capture of mice layer 2/3 visual cortex (**Figure 5)**, optical imaging containing astrocytic calcium activity within the spinal cord of free moving mice (45 Hz) (**Figure 6)**, and *ex vivo* (1.1 Hz) imaging of coronal, acute, neocortical slices (400 μm thick) in P10-23 mice (**Figure 4**). We demonstrated correct operation of all steps of the detection pipeline (**Figure 4, Figure 5**) with identically detected events corresponding to those identified by AQuA2. Furthermore, testing of AQuA2-Cloud across several trials of mouse spinal cord imaging yielded quantification of spatially-local calcium activity events in several regions of bulk glial cell population (**Figure 6**), matching those detected by AQuA2. For the *in vivo* test dataset, propagation metrics were identically computed for events derived from a split super-event (**Figure 5G-I**). While processing this dataset, implemented algorithm validity concerning spatial and temporal segmentation components of the detection pipeline were verified for correctness; temporal segmentation, noted to be reworked in [16], via a high sensitivity splitting of a super-event was apparent (**Figure 5E**) when contrasted against a (**Figure 5D**) pre-spatial segmentation reference. Comparison of compute time performance (**Table 1**) between AQuA2 and AQuA2-Cloud showed no discrepancy greater than 6.6%: attributable to natural system variations and overhead due to containerization.

**Figure 5:**
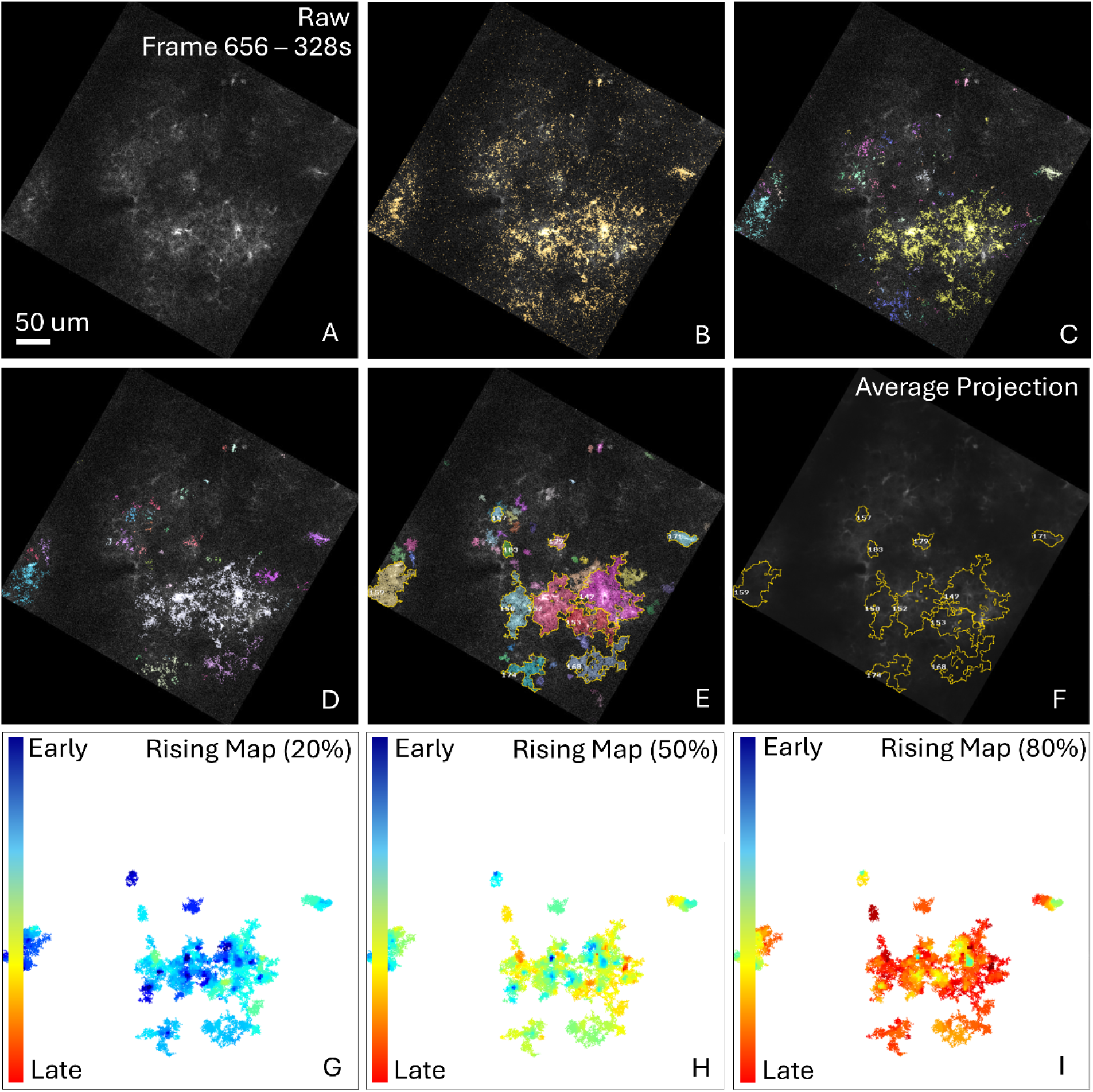
Detection pipeline step-wise results for *in vivo* data. Depicted is activity detection in imaging data containing astrocytic calcium expression with detection pipeline step-wise visual results (A-E). Raw imaging data (A) is analyzed for statistical significance of active voxels exhibiting potential signals (B) for which temporal segmentation is then performed to generate initial detected events (C). Spatial segmentation further refines the event population (D-E) prior to the user then being able to quantify selected events (E). Depicted example quantification includes event spatial projection (F) and event propagation characteristics (G-I).

**Figure 6:**
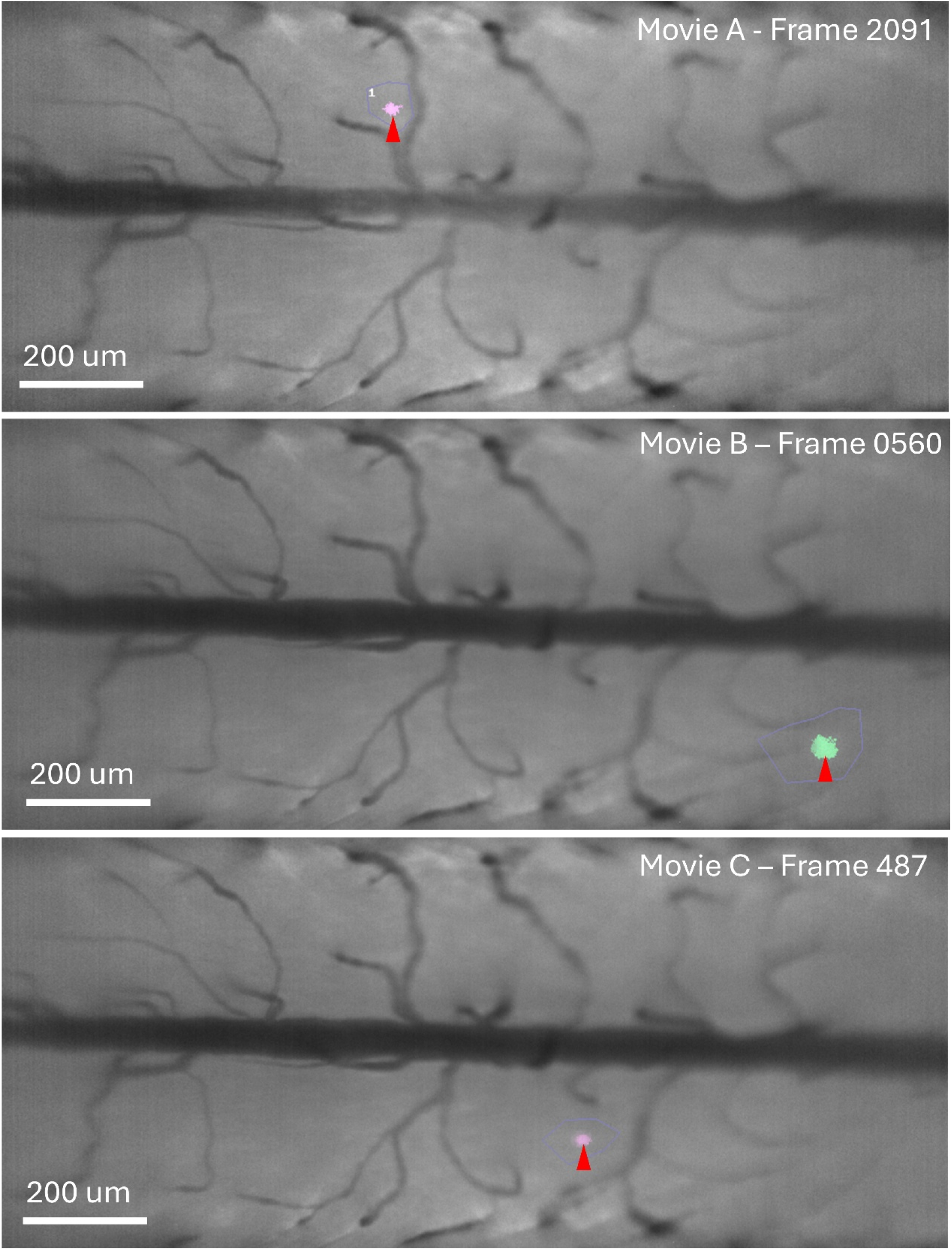
Small signal detection in mouse spinal cord imaging data. AQuA2-Cloud allows for detection of events within large imaging datasets. This includes global spatiotemporal signals as well as hidden local signals that may be components of larger global events. Depicted here are identified instances of spatially local calcium activity within trans-segmental optical capture of mouse spinal cord exhibited via GCaMP6..

**Table 1:** Compute time comparison between AQua2 and AQuA2-Cloud.

| Dataset | Data Size | AQuA2 | AQuA2-Cloud | Difference |
| --- | --- | --- | --- | --- |
| *Ex Vivo* Imaging Data | 502x502x281 | 81.4s | 81.2s | +0.2 % |
| *In Vivo* Imaging Data | 654x654x870 | 210.0s | 217.2s | -3.3% |
| Mouse Spinal Cord | 956x422x3122 | 33m 26.6s | 31m 13.4s | +6.6% |

## Conclusion

Compared to AQuA2, AQuA2-Cloud offers several advantages. AQuA2-Cloud offers access to the same analysis algorithms and processing pipeline found in AQuA2 in a centralized location without any end user licensing or environment restrictions. Any device with a web browser can utilize the cloud interface and conduct such analyses, and multiple users are able to process data simultaneously. The system is hosted in a portable docker container and only the deployer of the container needs to provide a MATLAB license and have some technical knowledge. The centralization of compute and storage capacity has benefits for user groups that may wish to share compute and storage resources with collaborators and others, or share results of analyses. Version control is provided in that all cloud users effectively utilize the same MATLAB installation and version as well as the same logic for analyses. Updating the AQuA2-Cloud version, and by extension the analysis logic utilized by all users, is a simple matter of obtaining the latest files from the public repository then redeploying the container. This may offer greater simplicity compared to systems with multiple MATLAB installations and duplicate copies of AQuA2 across multiple users. Integrated file management and referencing functions within the web interfaces, as well as a database connected FTP server, provide more intuitive mechanisms for uploading raw data and downloading analyses results compared to non-service-integrated file transfer systems. Files uploaded and results generated are immediately visible within the cloud application’s integrated file manager interface. Furthermore, project files generated by AQuA2-Cloud are directly and cross-platform compatible with AQuA2. Finally, the container and system employ internal CPU-based graphical rendering. Therefore, AQuA2-Cloud can be deployed on headless servers that do not have any GUI environments or libraries. This system allows investigators and end users with minimal technical knowledge the capability to conduct highly accurate and visually-interactive analysis of spatiotemporal activity within biological imaging movies with computations offloaded to a remote server.

## Methods

### Dataset: *Ex vivo* imaging

Coronal, acute neocortical slices (400 µm thick) from P10-23 mice were imaged using two-photon (2P) scanning at 502x502 pixel resolution. *Ex vivo* samples were imaged at 1.1 Hz frame rate (approx. 0.3 µm/pixel) with astrocytic GCaMP6 expression providing quantifiable fluorescence activity (**Figure 4**) within the optical field while slices were maintained in continuously aerated, standard artificial cerebrospinal fluid (ACSF). A single cell is shown with its primary activity detected, along with several spatially-local clustered events. No image registration or bleach correction was required. Data was processed using default pipeline settings.

### Dataset: *In vivo* imaging

*In vivo* 2 Hz fluorescence imaging data of visual cortex layers 2/3 at 512x512 pixel resolution (rotated within a 654x654 frame, 0.8 µm/pixel) of awake mice head-fixed on a circular treadmill. Astrocytic GCaMP6 fluorescence reveals Ca^2+^ activity across multiple astrocytes. AQuA2-Cloud quantified several events, including a super-event that was subsequently split (**Figure 5E**). No image registration or bleach correction was required. Processing pipeline was modified only by increasing *Source detection sensitivity* to 10 during spatial segmentation.

### Dataset: Mouse Spinal Cord

Lumbar spinal cord imaging data of freely moving GFAP-GCaMP6f mice at a pixel resolution of 956x422, 45 Hz frame rate, and 1.62 µm/pixel spatial scale. Mice were free to move about an enclosed space while their spinal cord were optically imaged [6]. The data were illumination corrected, motion-corrected, and cropped in time and space. A tail pinch was delivered once during the imaging sequence to evoke global calcium activity across the optical field; however, this global event is not shown, as the analysis focused on spatially-local activity. Three separate imaging movies from the dataset were evaluated, each demonstrating localized calcium transients indicated by small red arrows (**Figure 6**).

For the first movie (**Figure 6A**), preprocessing was performed and a cell boundary drawn as depicted. The *intensity threshold scaling* parameter was set to 1.2 and *maximum size* set to 500 pixels, with all other settings left at default values. For the second movie (**Figure 6B**), preprocessing and boundary definition were repeated with *intensity threshold scaling* set to 1.6 and *maximum size* to 600 pixels. For the third movie (**Figure 6C**), the same procedure was applied with *intensity threshold scaling* set to 2.1 and *maximum size* to 500 pixels. All three movies were processed at full spatial resolution to detect localized calcium activity, while the first movie was additionally downsampled by a factor of two then processed using fully default pipeline settings for performance evaluation (**Table 1**).

### Performance Evaluation

Platform compute time performance was compared between a standalone AQuA2 instance and an AQuA2-Cloud container instance, with both running inside virtual machines (VMs) configured to emulate the hardware of a typical compute server. For the AQuA2 standalone instance, local control within its VM was used to execute the full detection-pipeline analysis on a given dataset. For the AQuA2-Cloud instance, a remote user executed the same analysis workflow via the web interface. All runs were initialized only after both instances were fully deployed and the host and guest operating systems were idle. This procedure was repeated for all datasets across several trials.

Total compute time was defined as the duration between issuing the command to execute all detection-pipeline steps and the completion of the user-interface render and imaging-frame display following processing. Default pipeline settings were used for all datasets and trials. Both the AQuA2 VM and the deployed AQuA2-Cloud container instance were assigned identical CPU and RAM allocations.

### Workstation Configuration

The host for compute time analysis was a custom-built server equipped with an AMD EPYC 9684X processor (2.55 GHz base clock) and 384 GB of RAM. VMware Workstation 16.2.3 was used as the hypervisor for virtual machine (VM) hosting. The AQuA2 VM was allocated 24 CPU cores, 64 GB of RAM, and Ubuntu 24.04.2 as the operating system. The AQuA2-Cloud VM, which hosted the AQuA2-Cloud Docker container and its associated volume, was allocated 36 CPU cores, 96 GB of RAM, and the same operating system. The AQuA2-Cloud container itself was deployed with 24 CPU cores and allocated 64 GB of RAM to match the AQuA2 VM’s specifications. The C++ accelerated components of AQuA2 and the AQuA2-Cloud software stack were compiled utilizing the G++ 10 compiler. No GPUs were used in the performance evaluations.

## Availability and Requirements

Project Name: AQuA2-Cloud

Project Home Page: https://github.com/yu-lab-vt/AQuA2-Cloud

Operating System: Platform Independent

Programming Language: JavaScript, PHP, MATLAB (None needed by users)

Other Requirements: Single MATLAB license (For deployer)

License: GNU GPL, MIT

Any restrictions to use by non-academics: None

## Declarations

### Ethics approval and consent to participate

Not applicable

### Consent for publication

Not applicable

### Availability of data and materials

All testing data has been deposited at (https://data.mendeley.com/datasets/snkwr5bhnk/2).

### Author’s Contributions

MB (Conceptualization [Lead], Formal Analysis [Lead], Methodology [Lead], Software [Lead], Writing-original draft [Lead], Writing-review & editing [Lead]), XM (Software [Supporting]), DD (Data curation [Equal], Writing-review & editing [Supporting]), EC (Data curation [Equal], Writing-review & editing [Supporting]), BL (Software [Supporting], Writing-review & editing [Supporting]), YW (Software [Supporting], Resources [Supporting]), AN (Data curation [Supporting]), GY (Supervision [Lead], Funding acquisition [Lead], Methodology [Supporting], Writing-review & editing [Supporting])

## Acknowledgments

Not applicable

## References

1. Feng J, Zhang C, Lischinsky JE, Jing M, Zhou J, Wang H, Zhang Y, Dong A, Wu Z, Wu H, Chen W. A genetically encoded fluorescent sensor for rapid and specific in vivo detection of norepinephrine. Neuron. 2019 May 22;102(4):745–61.

2. Wu Z, He K, Chen Y, Li H, Pan S, Li B, Liu T, Xi F, Deng F, Wang H, Du J. A sensitive GRAB sensor for detecting extracellular ATP in vitro and in vivo. Neuron. 2022 Mar 2;110(5):770–82.

3. Masharina A, Reymond L, Maurel D, Umezawa K, Johnsson K. A fluorescent sensor for GABA and synthetic GABAB receptor ligands. Journal of the American Chemical Society. 2012 Nov 21;134(46):19026–34.

4. Dong C, Zheng Y, Long-Iyer K, Wright EC, Li Y, Tian L. Fluorescence imaging of neural activity, neurochemical dynamics, and drug-specific receptor conformation with genetically encoded sensors. Annual review of neuroscience. 2022 Jul 8;45:273–94.

5. Bear M, Connors B, Paradiso MA. Neuroscience: Exploring the brain. Jones & Bartlett Learning; 2025 Jul 11.

6. Shekhtmeyster P, Duarte D, Carey EM, Ngo A, Gao G, Olmstead JA, Nelson NA, Nimmerjahn A. Trans-segmental imaging in the spinal cord of behaving mice. Nature Biotechnology. 2023 Dec;41(12):1729–33.

7. Shekhtmeyster P, Carey EM, Duarte D, Ngo A, Gao G, Nelson NA, Clark CL, Nimmerjahn A. Multiplex translaminar imaging in the spinal cord of behaving mice. Nature Communications. 2023 Mar 21;14(1):1427.

8. Patriarchi T, Cho JR, Merten K, Howe MW, Marley A, Xiong WH, Folk RW, Broussard GJ, Liang R, Jang MJ, Zhong H. Ultrafast neuronal imaging of dopamine dynamics with designed genetically encoded sensors. Science. 2018 Jun 29;360(6396):eaat4422.

9. Jing M, Zhang P, Wang G, Feng J, Mesik L, Zeng J, Jiang H, Wang S, Looby JC, Guagliardo NA, Langma LW. A genetically encoded fluorescent acetylcholine indicator for in vitro and in vivo studies. Nature biotechnology. 2018 Aug;36(8):726–37.

10. Giovannucci A, Friedrich J, Gunn P, Kalfon J, Brown BL, Koay SA, Taxidis J, Najafi F, Gauthier JL, Zhou P, Khakh BS. CaImAn an open source tool for scalable calcium imaging data analysis. elife. 2019 Jan 17;8:e38173.

11. Pachitariu M, Stringer C, Schröder S, Dipoppa M, Rossi LF, Carandini M, Harris KD. Suite2p: beyond 10,000 neurons with standard two-photon microscopy. BioRxiv. 2016 Jun 30:061507.

12. Wang Y, DelRosso NV, Vaidyanathan TV, Cahill MK, Reitman ME, Pittolo S, Mi X, Yu G, Poskanzer KE. Accurate quantification of astrocyte and neurotransmitter fluorescence dynamics for single-cell and population-level physiology. Nature neuroscience. 2019 Nov;22(11):1936–44.

13. Kustikova V, Krivonosov M, Pimashkin A, Denisov P, Zaikin A, Ivanchenko M, Meyerov I, Semyanov A. CalciumCV: Computer vision software for calcium signaling in astrocytes. InInternational Conference on Analysis of Images, Social Networks and Texts 2018 Jul 5 (pp. 168–179). Cham: Springer International Publishing.

14. Bjørnstad DM, Åbjørsbråten KS, Hennestad E, Cunen C, Hermansen GH, Bojarskaite L, Pettersen KH, Vervaeke K, Enger R. Begonia—a two-photon imaging analysis pipeline for astrocytic ca2+ signals. Frontiers in Cellular Neuroscience. 2021 May 20;15:681066.

15. Renton AI, Dao TT, Johnstone T, Civier O, Sullivan RP, White DJ, Lyons P, Slade BM, Abbott DF, Amos TJ, Bollmann S. Neurodesk: an accessible, flexible and portable data analysis environment for reproducible neuroimaging. Nature methods. 2024 May;21(5):804–8.

16. Mi X, Chen AB, Duarte D, Carey E, Taylor CR, Braaker PN, Bright M, Almeida RG, Lim JX, Ruetten VM, Wang Y. Fast, accurate, and versatile data analysis platform for the quantification of molecular spatiotemporal signals. Cell. 2025 May 15;188(10):2794–809.

17. Mi X, Wang M, Chen A, Lim JX, Wang Y, Ahrens MB, Yu G. BILCO: An Efficient Algorithm for Joint Alignment of Time Series. Advances in Neural Information Processing Systems. 2022 Dec 6;35:36270–81.

18. Rohnert H. {Pattern-oriented} Software architecture. In2nd USENIX Conference on Object-Oriented Technologies (COOTS 96) 1996.

19. Bass L. Software architecture in practice. Pearson Education India; 2012.

20. Gabarró S. Web application design and implementation: Apache 2, PHP5, MySQL, JavaScript, and Linux/UNIX. IEEE Computer Society; 2007 Mar 23.

21. Kenyon T. Data networks: routing, security, and performance optimization. Elsevier; 2002 Jul 18.

22. Shyam G. Cloud computing: Concepts and technologies. CRC Press; 2021 Mar 8.

